# Loss of 5-HT_2C_ Receptor Function Alters Motor Behavior in Male and Female Mice With and Without Spinal Cord Injury

**DOI:** 10.1101/2025.02.06.636971

**Authors:** Margaret I. Sim, Derin Birch, Amr A. Mahrous, C.J. Heckman, Vicki M. Tysseling

## Abstract

The 5-HT_2C_ receptor is involved in the regulation of spinal motor function, specifically in both volitional and involuntary motor behavior. It contributes to various aspects of voluntary movement, such as locomotion, gait, coordination, and muscle contraction, as well as to involuntary motor behavior like spasms, which affect many individuals with spinal cord injury. Despite its known involvement in motor function, little is known about the physiological role of the 5-HT_2C_ receptor and the changes it undergoes after spinal cord injury. In this study, we have investigated the volitional and involuntary motor behavior of male and female uninjured and spinal cord injured knock-out mice that lack the functional 5-HT_2C_ receptor by comparing these genetically manipulated mice to typical-functioning sex-matched wildtype mice. Behavioral assessments revealed differences in volitional muscle strength and coordination, as well as hyperreflexia, between the groups observed. Additionally, *ex vivo* sacral cord preparation data suggest that 5-HT_2C_ receptor knock-out mice exhibit less spasm-like activity than wildtype mice, corroborating our results from behavioral testing in which the flexor withdrawal reflex of the hindlimbs was assessed. To investigate potential compensatory changes in 5-HT_2C_ receptor expression following spinal cord injury, western blot analysis was performed on lumbar and sacral spinal cord tissue from wildtype and 5-HT_2C_ receptor knock-out mice before and after injury. Both sex and injury status significantly influenced 5-HT_2C_ receptor relative expression and distribution of these receptors in both spinal cord regions. Through a comprehensive approach combining behavioral assessments, electrophysiological experiments, and whole-tissue protein analysis, our findings provide strong evidence that the 5-HT_2C_ receptor plays a critical role in both volitional motor function and involuntary motor behavior, and the relative expression of the 5-HT_2C_ receptor is influenced by both sex and injury status.

## Introduction

Serotonin (5-HT) is a complex neuromodulator that has been implicated in a variety of physiological functions (Mosher et al., 2005). In motor control, descending 5-HT within the spinal cord is responsible for the regulation of spinal motoneuron (MN) excitability (Bayliss et al., 1995; Heckman et al., 2003; Lee and Heckman, 1998; Lindsay and Feldman, 1993; Zhang et al., 1997) through the activation of 5-HT_2_ receptors. The 5-HT_2_ class of receptors are G-protein coupled receptors (GPCRs), of which there are three subfamilies: the 5-HT_2A_, 5-HT_2B_, and 5-HT_2C_ receptors (Barnes and Sharp, 1999). All three of the 5-HT_2_ receptor subtypes have been shown to possess similar molecular structure, pharmacology, and signal transduction pathways to one another (Parajulee and Kim, 2023). Of the three 5-HT_2_ receptor subtypes, 5-HT_2A_ and 5-HT_2C_ have been the most extensively studied for their role in motor function.

Motoneurons express voltage-gated Cav1.3 L-type calcium channels which mediate persistent inward currents (PICs) (Binder et al., 1996). These channels can amplify synaptic inputs and generate sustained depolarization in response to brief excitatory inputs (Bennett et al., 2004; Heckman et al., 2005). The presence of serotonin and the activation of the 5-HT_2_ receptors have been shown to be essential for facilitating PICs in the intact as well as the injured spinal cord (Hounsgaard et al., 1988; Murray et al., 2010, 2011b). Despite these potent effects that 5-HT has on MN excitability, the role of different 5-HT_2_ receptors in motor behavior is still not fully understood.

Previous studies have shown that pharmacologically blocking the 5-HT_2C_ receptor (5-HT_2C_R) has no effect on locomotion or reflexes (Majczyński et al., 2020). However, several off-target effects of the pharmacological blockers might have affected this outcome. More recent research using knock-out (KO) 5-HT_2C_R mice have shown conflicting reports of the effect on locomotion in intact animals (Heisler et al., 2007; Hill et al., 2011; Nebuka et al., 2020), highlighting a need for further investigation. Additionally, the 5-HT_2C_R has been shown to be involved in the development of muscle spasms following spinal cord injury (SCI) (Murray et al., 2010; Tysseling et al., 2017), suggesting potential therapeutic interventions through manipulating these receptors. Furthermore, it is unknown whether these receptors mediate sex differences in normal motor behaviors or motor recovery after SCI given the reported relationship between sex hormones (i.e. estradiol and testosterone) and serotonin receptor expression (Kugaya et al., 2003; Popova and Amstislavskaya, 2002). Therefore, the goal of the current study is to investigate the role of the 5-HT_2C_R in motor behavior in uninjured and spinal cord injured wildtype (WT) and genetically-modified KO mice that lack the functional 5-HT_2C_R (5-HT_2C_R KO).

Recent evidence has shown differences in PIC generation between male and female human subjects (Jenz et al., 2023). This study found that although motor unit discharge rates did not differ between male and female healthy subjects, biological sex was a critical variable in estimating PIC magnitude. Female subjects were revealed to have larger PIC estimates in lower limb MNs than their male counterparts. In light of these findings, we deemed it necessary to include sex-specific analyses across all experimental procedures conducted.

The second purpose of this study was to investigate potential differences in involuntary motor behavior post-SCI between the 5-HT_2C_R KO mice and their WT counterparts. This study is unique because a thoracic model of injury is used, whereas previous research has typically used a sacral transection, and involuntary motor behavior has been examined only in the sacrocaudal spinal cord. It is well understood that injury severity and recovery outcome are predominately influenced by the level of SCI. Using a thoracic model of spinal cord transection is more clinically relevant, as approximately 35% of all SCIs occur at the thoracic level (Alizadeh et al., 2019), and it allows us to examine mechanistic alterations in the lumbar spinal cord, as the descending pathways that project to the lumbar section are responsible for hindlimb locomotion (Han et al., 2019).

To further investigate mechanistic alterations underlying involuntary motor behavior post-SCI, we analyzed total protein expression in the lumbar and sacral spinal cord. Whether the 5-HT_2C_R is upregulated post-SCI remains unclear, as total protein and mRNA studies have yielded conflicting results. While total mRNA studies in rats suggest that the 5-HT_2C_R is not upregulated after sacral transection, constitutive activity of the receptor has been shown to increase post-SCI (Murray et al., 2010). Conversely, immunoreactivity and total protein analyses indicate that the 5-HT_2C_R is upregulated in the sacral cord 60 days post-sacral transection (Murray et al., 2010; Ren et al., 2013). However, it is still unknown whether the distribution of the 5-HT_2C_R is altered in the lumbar and sacral spinal cord following a thoracic spinal cord transection. In line with the second objective of this study, we aim to clarify discrepancies in lumbar and sacral protein expression to better understand the underlying molecular adaptations contributing to the facilitation of involuntary motor behavior post-SCI. The results of this study suggest the possibility of there being a homeostatic relationship or feedback loop that is often characterized by GPCRs.

## Materials and Methods

### Animals

All procedures in this study were reviewed and approved by the Northwestern University Institutional Animal Care and Use Committee (IACUC) and were compliant with the National Institutes of Health Guide for the Care and Use of Laboratory Animals. Two groups of adult mice (Jackson Laboratory, Bar Harbor, ME, USA) inclusive of both sexes were used in this study.

Transgenic knockout mice, B6.129-Htr2c^tm1Jul/J^ (5-HT_2C_R KO), which lack the functional serotonin 2C receptor, were compared with age-matched wild-type C57Bl/6J mice (WT). Following behavioral testing in intact animals, both experimental groups received spinal cord injury at 10-weeks of age and were compared to uninjured littermate controls. Behavioral testing was performed before and after SCI and was followed by terminal procedures in which either proteins were extracted for western blot analysis or the sacrocaudal spinal cord was removed for *ex vivo* analysis (see below).

### Spinal Cord Injury

At ten weeks of age, mice were anesthetized with isoflurane, and a laminectomy was performed at the T10 spinal level to expose the T11-T12 spinal segments. The spinal cord was completely transected using spring scissors. Following transection, mice were evaluated for 10 weeks using the Basso Mouse Scale (see below) to measure functional motor recovery. SCI was characterized as ‘chronic’ 10 weeks post-surgery. At this point, the test subjects underwent more behavioral testing before the terminal procedure of either protein extraction or sacrocaudal spinal cord removal.

### Volitional Motor Behavior Testing

#### Basso Mouse Scale

The Basso Mouse Scale (BMS) is an established open-field behavioral assessment used to evaluate hindlimb functional deficits following spinal cord injury in mice (Basso et al., 2006). 5-HT_2C_R KO (n = 11) and WT (n = 15) SCI mice underwent weekly evaluations beginning one day post-SCI (week 0) and continuing through until the injury was chronic (week 10). The scoring of this assessment ranges from 0, indicating complete paralysis and total loss of hindlimb function, to 9, indicating standard functionality and locomotive ability. Sub-scales were not necessary to include due to low function. An average of the left and right hindlimb scores were taken to obtain the BMS score for each subject. All test subjects retained a BMS score of < 2 throughout the experiment.

#### DigiGait™

Treadmill gait analysis was conducted using the DigiGait™ system (Mouse Specifics Inc., Framingham, MA) to assess specific differences in gait between uninjured 5-HT_2C_R KO (n = 48) and WT adult mice (n = 10). The protocol for this assessment has been discussed previously (Mancuso et al., 2011). Briefly, the DigiGait™ system consisted of a transparent treadmill fixed horizontally at 0° (5 cm in width, 25 cm in length) that restricted the mice as they ran at a constant velocity. The velocity used in this assessment was fixed at 24 cm/s, a speed that mice maintained without veering off to one side. A high-temporal-resolution video recording (∼5 seconds) was captured for each mouse before analysis using the DigiGait™ Imaging and Analysis software v12.2 (Mouse Specifics Inc., Framingham, MA). The software utilizes coding and artificial intelligence algorithms to detect the spatial coordinates and direction of pixels, enabling it to calculate either the duration spent in each gait phase or the distance between each paw. Data was averaged between left and right paws in both the forelimbs and hindlimbs.

#### Grip Strength

A grip strength meter (Ugo Basile, Gemonio, Italy) was utilized to determine the maximum and average force displayed by the mice in the forelimbs, hindlimbs, and all four limbs combined. This experimental procedure has been described previously (Papaneophytou et al., 2018).

Uninjured male 5-HT_2C_R KO (n = 12) and WT (n = 18) mice and female 5-HT_2C_R KO (n = 14) and WT (n = 19) mice were assessed. For forelimb and all-limb testing, each mouse was held by the base of the tail and lowered onto the plastic grid to grip with its forepaws or all four paws.

The mouse was then pulled away slowly, with the torso horizontal to the table, allowing the limbs to flex until the grip was released (Figure 4A). For hindlimb testing, the mice gripped onto the triangle bar attachment with their forelimbs and were then lowered down until only the hindpaws were able to grip onto the plastic attachment. The same parallel pulling motion was used. A digital force transducer recorded the peak pull force (g). Tension was recorded when the mouse released the grip from the attachment. Each mouse underwent five trials per grip type, with about 30 minutes between each trial. The order in which the mice were tested was randomized, and the experimenter was blinded to both the strain and sex of each mouse that was assessed.

### Involuntary Motor Behavior Testing

#### Flexor Withdrawal Testing

The flexion reflex analysis was conducted on 5-HT_2C_R KO (n = 11) and WT (n = 14) SCI mice to evaluate hyperreflexia and muscle spasms in the hindlimb. This experimental procedure has been described in detail previously (Tysseling et al., 2017). Briefly, electromyography (EMG) recording electrodes were placed in the tibialis anterior (TA) and lateral gastrocnemius (LG) muscles of the hindlimb under isoflurane anesthesia. Ball electrodes that were used to deliver electrical stimulations were then affixed to the plantar side of the hindpaw to evoke the flexion withdrawal reflex. When the mouse was fully awake following cessation of gaseous anesthesia, the response threshold was determined by increasing the amplitude of current by 10 µA increments until an observable movement in the hindlimb could be identified. After the threshold had been determined, 10 trains (5 pulses, 1 ms pulse width, 100 Hz) were applied at 5x threshold with a minimum of 2 minutes of rest between stimuli, until a total of 5 viable recordings were obtained. A trial was considered viable when there was no substantial EMG activity for at least 1 second before the stimulation was applied. The final score was presented as the integral of the rectified EMG response normalized to baseline, with a magnitude of 0 indicating that no response was calculated above the baseline. The evoked reflex was divided into the short polysynaptic reflex (SPR; 0 ms – 40 ms), longer polysynaptic reflex (LPR; 40 ms – 500 ms), the long-lasting reflex (LLR; subdivided into LLR1, 500 ms – 1.5 s, LLR2, 1.5 s – 2.5 s, and LLR 3, 2.5 s – 3.5 s), the total long-lasting reflex (total LLR, 500 ms – 3.5 s), and the total signal (summed trace; 40 ms – 3.5 s).

#### Ex Vivo Sacral Cord Preparation

The detailed methodology for this experiment has been previously described (Mahrous and Elbasiouny, 2017). Briefly, mice were anesthetized with an intraperitoneal injection of urethane (≥ 0.2 g/100g). A dorsal laminectomy exposed the lower half of the spinal cord, and a transection at L5-L6 enabled the removal of the entire sacrocaudal spinal cord (S1-Co2 segments) along with its attached spinal roots. The extracted tissue was maintained *ex vivo* in oxygenated artificial cerebrospinal fluid (ACSF) at room temperature (∼21°C) and allowed to rest for ∼ 1 hour before any stimulation. The dorsal roots (DRs) and ventral roots (VRs) on both sides of the cord were mounted on bipolar wire electrodes. DRs were connected to an S88 Grass stimulator (A-M Systems), while VRs were connected to differential amplifiers (WPI, 1000xgain, filtered between 300 Hx-3kHz). Trains of stimulation were delivered to the DRs (5 pulses, 0.1 ms pulse width, 25 Hz) at intensities of 2xT and 10xT, where T represents the threshold – the minimal stimulation amplitude required to elicit a detectable VR response. The evoked VR responses were quantified by measuring the area under the curve of the integrated signal. To block synaptic inhibition, strychnine and picrotoxin (STR/PTX) were added to the ACSF, and ∼ 15 minutes were allowed to achieve full drug effects. Following drug application, DR stimulation elicited VR responses that persisted for several seconds (spasm-like activity). To prevent adaptation of spasm-like activity with repeated stimulations (Mahrous et al., 2024), a minimum 60-second period without pre-stimulation activity was ensured. The spasm-like activity was analyzed by dividing the evoked VR signal into the short polysynaptic reflex (SPR; 0 ms – 40 ms, data not shown), longer polysynaptic reflex (LPR; 40 ms – 500 ms), the long-lasting reflex (LLR; subdivided into LLR1, 500 ms – 1.5 s, LLR2, 1.5 s – 2.5 s, and LLR 3, 2.5 s – 3.5 s), the total long-lasting reflex (total LLR, 500 ms – 3.5 s), and the total signal (summed trace; 40 ms – 3.5 s).

### Protein Analysis

#### Tissue Preparation

Uninjured male 5-HT_2C_R KO (n = 4) and WT (n = 4) mice and female 5-HT_2C_R KO (n = 4) and WT (n = 4) mice, as well as injured male 5-HT_2C_R KO (n = 4) and WT (n = 4) mice and female 5-HT_2C_R KO (n = 4) and WT (n = 4) mice were used for western blot analysis. Spinal cord tissue from the lumbar and sacral section of the cord was identified and separated beneath the ventral root L6. The tissue was immediately placed on ice and prepared for homogenization by sonication with T-PER (Thermo Scientific; Waltham, MA, USA) and Halt Protease Inhibitor Cocktail (Thermo Scientific; Waltham, MA, USA) and centrifuged at 10,000-g for 5-mins.

Supernatant was separated and aliquoted prior to being stored at -80°C until ready for use.

#### Western Blot

The protein concentration of tissue supernatant was determined using Pierce BCA Protein Assay reagent kit (Thermo Scientific; Waltham, MA, USA) in order to ensure equal protein loading per lane. Sample preparation consisted of combining the lysed and homogenized sample with sample buffer and reducing agent (BioRad; Hercules, CA, USA), and T-PER before boiling at 100°C for 5-mins. For each sample, 12 µg of protein extract was separated by 10-15% SDS-PAGE (BioRad; Hercules, CA, USA) and transferred to PVDF membrane (Thermo Scientific; Waltham, MA, USA). Membranes were blocked in 5% non-fat dried milk solution for 1-hr at room temperature before overnight incubation at 4°C with antibodies against 5-HT_2C_R (Catalog #MA5-3271, 1:1,000, Thermo Scientific; Waltham, MA, USA), GAPDH (Catalog #MA5-35235, 1:100,000, Thermo Scientific; Waltham, MA, USA), 5-HT_2A_R (sc-166775, 1:500, Santa Cruz Biotechnology, TX, USA), and GAPDH (Catalog #AM4300, 3µg/mL, Thermo Scientific; Waltham, MA, USA). Following primary antibody incubation, each membrane was washed in TBS-T 3X for 5-mins/wash before secondary incubation with goat anti-rabbit (Catalog #31460, 1:1,000, Thermo Scientific; Waltham, MA, USA) or goat anti-mouse (Catalog #A-10668, 1:2,000, Thermo Scientific; Waltham, MA, USA) secondary antibody for 1-hr at room temperature. A chemiluminescence kit (SKU #AC2103, Azure Biosystems; Dublin, CA, USA) was used for immunoreactive band imaging using the Azure 400 system (AZI400-01, Azure Biosystems; Dublin, CA, USA). The western blots were analyzed using ImageJ software (FIJI) to quantify band intensities.

#### Statistical Analysis

Statistical analysis was done using GraphPad Prism Software (Version 10.0.1, GraphPad, Boston, MA) and IBM SPSS (Version 29.0.2.0, IBM Corp., Armonk, NY). Prior to statistical analysis, normality of the data was first assessed by using the Shapiro-Wilk test to determine whether the residuals follow a normal distribution. Diagnostic plots were generated to assess the assumptions of the two-way ANOVA. Residual plots, Q-Q plots, and homoscedasticity plots were examined to evaluate the assumptions of normality and homoscedasticity. If data passed normality as assessed by the Shapiro-Wilk test and the diagnostic plots showed no evidence of violations of homoscedasticity, the data was then evaluated using a two-way ANOVA without transformation. A parametric t-test was employed to analyze BMS score, while a two-way ANOVA was utilized for comparison of male and female, 5-HT_2C_R KO and WT mouse data from grip strength, DigiGait™, flexor withdrawal testing, sacral cord preparation, and western blot band intensity. A one-way ANOVA was used for burst duration analysis. Statistical significance is denoted as * p < 0.05, ** p < 0.01, *** p < 0.001, **** p < 0.0001.

## Results

### 5-HT_2C_R KO Mice Exhibit Decreased Power and Stability During Normal Locomotion

Previous experiments have described conflicting reports on the involvement of the 5-HT_2C_R in normal locomotion (Heisler et al., 2007; Hill et al., 2011). Here, we used the comprehensive DigiGait™ digital treadmill analysis to assess various gait parameters and identify subtle differences and possible motor impairments in both sexes of WT and 5-HT_2C_R KO mice during straight-line walking at a fixed velocity (see methods).

The stride phase consists of the swing (paw is in the air) and stance phases, with stance further divided into brake (heel contact) and propulsion (toe-off). To assess stride composition differences, the percentage of time spent in swing, brake, propel, and the combined brake and propel (stance) phases was analyzed in both the forelimbs and hindlimbs. Figure 1A presents forelimb data, showing that male 5-HT_2C_R KO and female WT mice spent less time in swing (*P* = 0.0347 and *P* = 0.0393, respectively) and more time in brake (*P* < 0.0001 and *P* = 0.0040, respectively) than male WT mice. Differences in the stance phase composition were sex- and genotype-dependent, with female (both WT and 5-HT_2C_R KO) mice spending a greater percentage of stride in stance compared to their male counterparts (*P* = 0.0393 and *P* = 0.0194, respectively). Additionally, both male and female 5-HT_2C_R KO mice spent a greater percentage of stride in the stance phase compared to their WT counterparts (*P* = 0.0347 and *P* = 0.0040, respectively).

**Figure 1.**
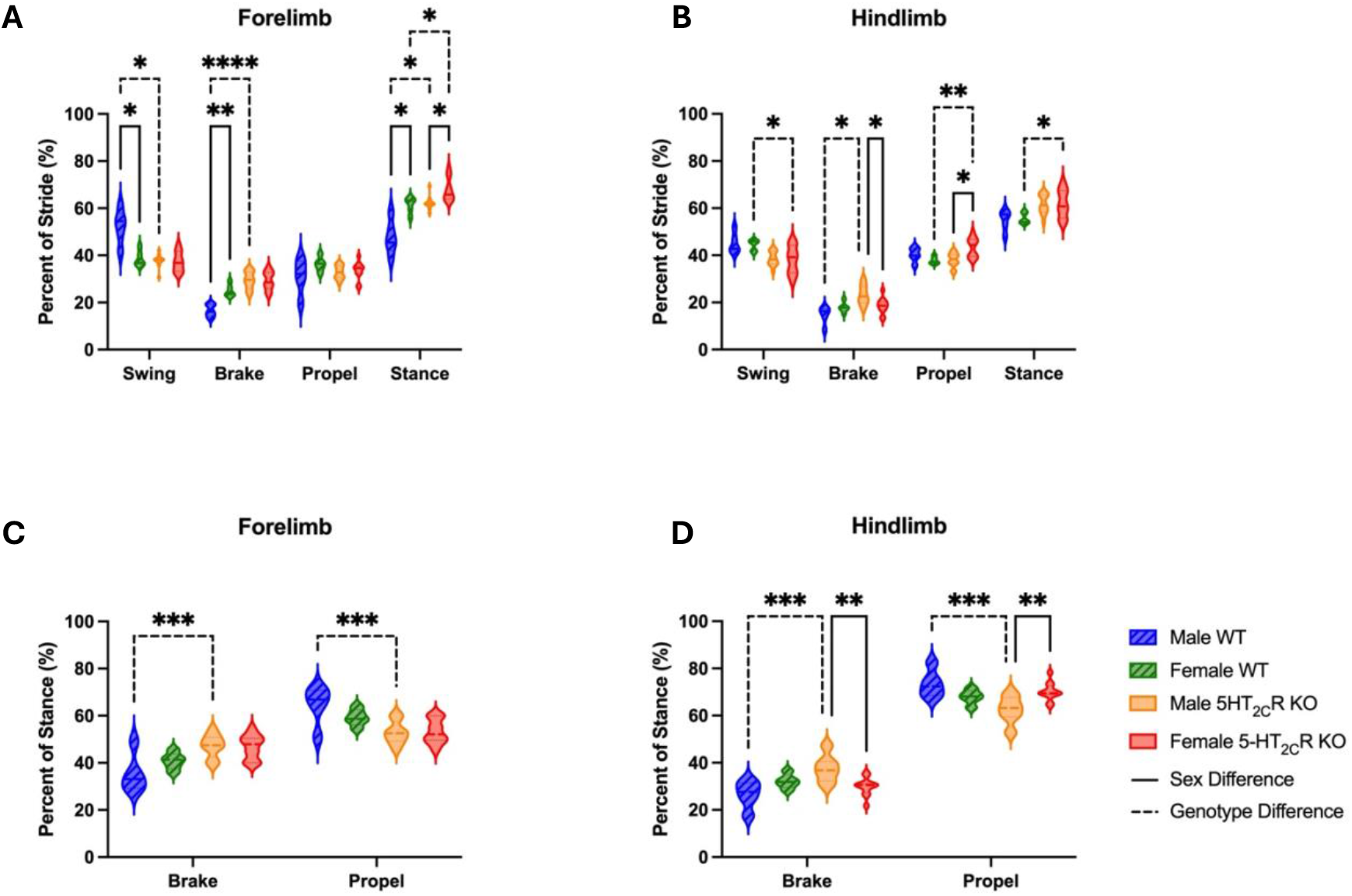
5-HT_2C_R KO mice have different gait phase composition than WT mice. A two-way ANOVA revealed significant differences in the forelimb and hindlimb composition of stride (inclusive of the swing, brake, propel, and stance (brake + propel) phases). Male WT (n = 5), female WT (n = 5), male 5-HT_2C_R KO (n = 12), and female 5-HT_2C_R KO (n = 11) mice were used in analysis. The p-value for significant differences is denoted in the figure above, with a solid significance line indicating a sex-specific difference and a dashed significance line indicating a genotype-specific difference. **** p < 0.0001, *** p < 0.001, ** p < 0.01, * p < 0.05.

Figure 1B shows the stride composition results from the hindlimbs, showing that female WT mice spent a greater percentage of stride in the swing phase than female 5-HT_2C_R KO mice (*P* = 0.0428). Differences in the brake and propel phase composition were sex- and genotype-dependent, as male 5-HT_2C_R KO mice spent a greater percentage of stride in brake (*P* = 0.0323) and less time in propel (*P* = 0.0130) than female 5-HT_2C_R KO mice. Additionally, male 5-HT_2C_R KO mice spent a greater percentage of stride in brake than male WT mice (*P* = 0.0127), and female 5-HT_2C_R KO mice spent a greater percentage of stride in propel compared to female WT mice (*P* = 0.0079). With the combined analysis of the brake and propel phase, it was revealed that female 5-HT_2C_R KO mice spent a greater percentage of stride in stance compared to female WT mice (*P* = 0.0428).

As previously mentioned, stance can be further divided into brake and propel. Upon independent examination of the stance phase, the forelimb analysis revealed that male 5-HT_2C_R KO mice spent a greater percentage of stance phase in propel and less in brake compared to male WT mice (*P* = 0.0007; Figure 1C). Additionally, hindlimb analyses revealed that both male WT and female 5-HT_2C_R KO mice spent a greater percentage of hindlimb stance in propel and less in brake than male 5-HT_2C_R KO mice (*P* = 0.0004 and *P* = 0.0019, respectively; Figure D), suggesting that both sex and genotype influenced the gait phase composition of stance.

Individual gait parameters were also examined. Stride length was assessed to investigate whether the sex or genotype of the mouse affected gait coordination and neuromuscular function as evaluated by the distance covered in taking one full stride. No difference in the stride length or the stride length coefficient of variance (measurement of variability within each group) was revealed (Figure 2A – B). After noting significant differences in the composition of stride and stance that suggest that 5-HT_2C_R KO mice may exhibit decreased stability compared to WT mice, we examined the stance width of each mouse group to illuminate potential differences in the distribution of weight in the forelimbs and hindlimbs. Figure 2C shows that female 5-HT_2C_R KO mice had a narrower stance width in both the forelimbs (*P* = 0.0272) and hindlimbs (*P* = 0.0064) compared to male 5-HT_2C_R KO mice, whereas only the hindlimb stance width of female WT mice was narrower than male WT mice (*P* = 0.0088). The variability of stance width in the forelimbs and hindlimbs of each mouse group was also evaluated using the coefficient of variance statistical measurement; however, significance was not attained in any group (Figure 2D). Notably, these results suggest an inherent difference in stance width between male and female mice, regardless of whether the 5-HT_2C_R is present.

**Figure 2.**
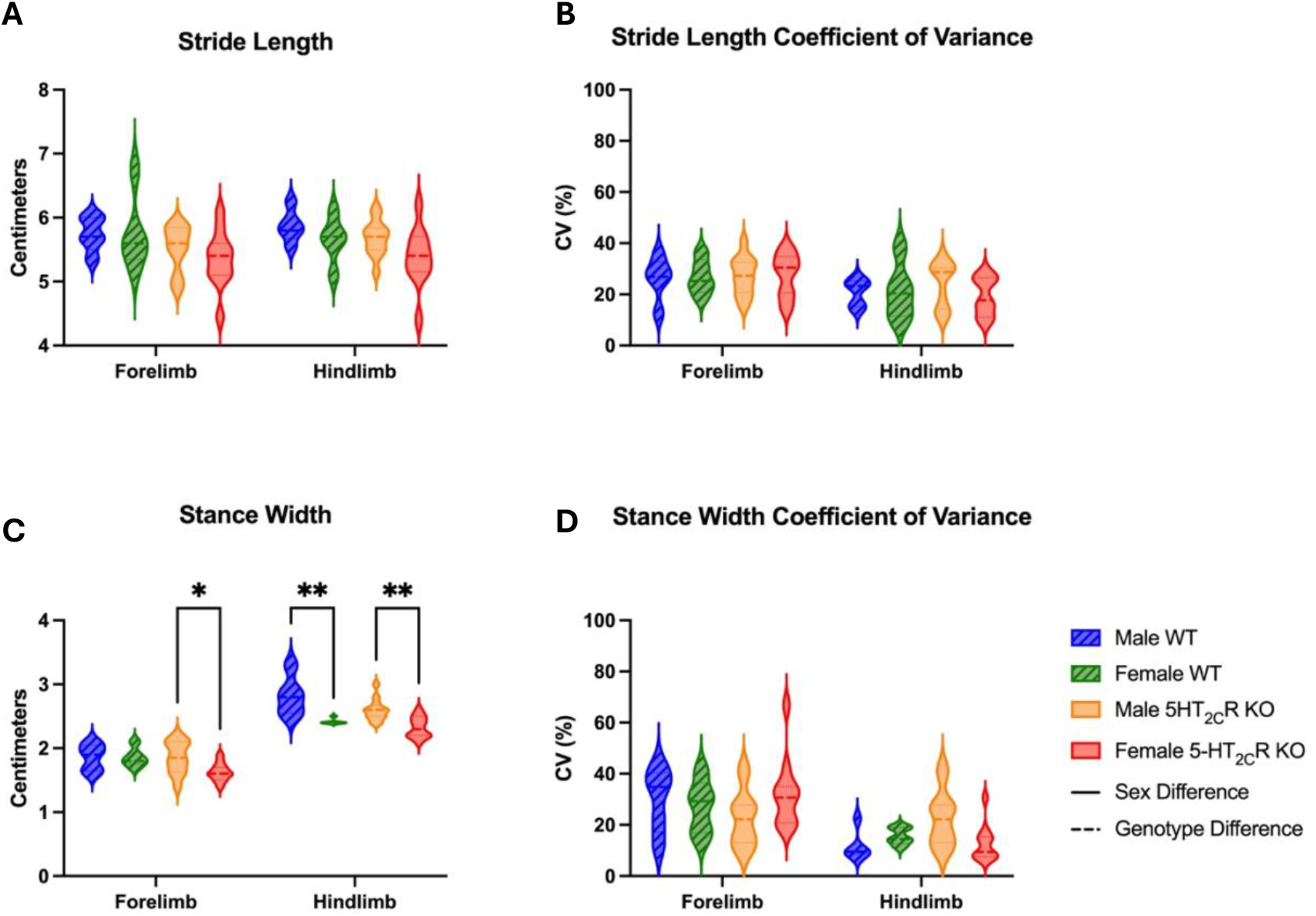
Stance width is affected by the sex of the mouse, regardless of the 5-HT_2C_R. A two-way ANOVA revealed significant differences in stance width between the forelimbs and hindlimbs, but not in stride length. Male WT (n = 5), female WT (n = 5), male 5-HT_2C_R KO (n = 12), and female 5-HT_2C_R KO (n = 11) mice were used in this analysis. The p-value for significant differences is denoted in the figure above, with a solid significance line indicating a sex-specific difference. **** p < 0.0001, *** p < 0.001, ** p < 0.01, * p < 0.05.

Following stance width analysis, we examined the area of the paw to possibly uncover differences in surface area and paw contact on the treadmill during normal locomotion. There were no differences in forelimb paw area between male or female WT and 5-HT_2C_R KO mice. However, the hindlimb paw area of both the female WT and male 5-HT_2C_R KO mice was significantly smaller compared to male WT mice (*P* = 0.0005 and *P* = 0.0089, respectively; Figure 3A), suggesting that the 5-HT_2C_R affects both the stability and weightbearing ability of male mice. Finally, midline distance was the last parameter assessed using the DigiGait™ system. Midline distance is the measurement of the total distance from the midline of the mouse’s body to its forelimb or hindlimb paw. This data provides insight into paw positioning relative to the midline, which can reflect the mouse’s relative stability. Both male WT mice and female WT mice had significantly larger forelimb midline distance compared to male 5-HT_2C_R KO mice (*P* = 0.0003) and female 5-HT_2C_R KO mice (*P* = 0.0225), however no difference in the hindlimb midline distance in any group was revealed (Figure 3B). These results suggest that the 5-HT_2C_R KO mice lack the necessary muscular power and strength to extend the forelimbs to the extent that the WT mice are capable of, thereby decreasing the foundational base of stability during normal locomotion.

**Figure 3.**
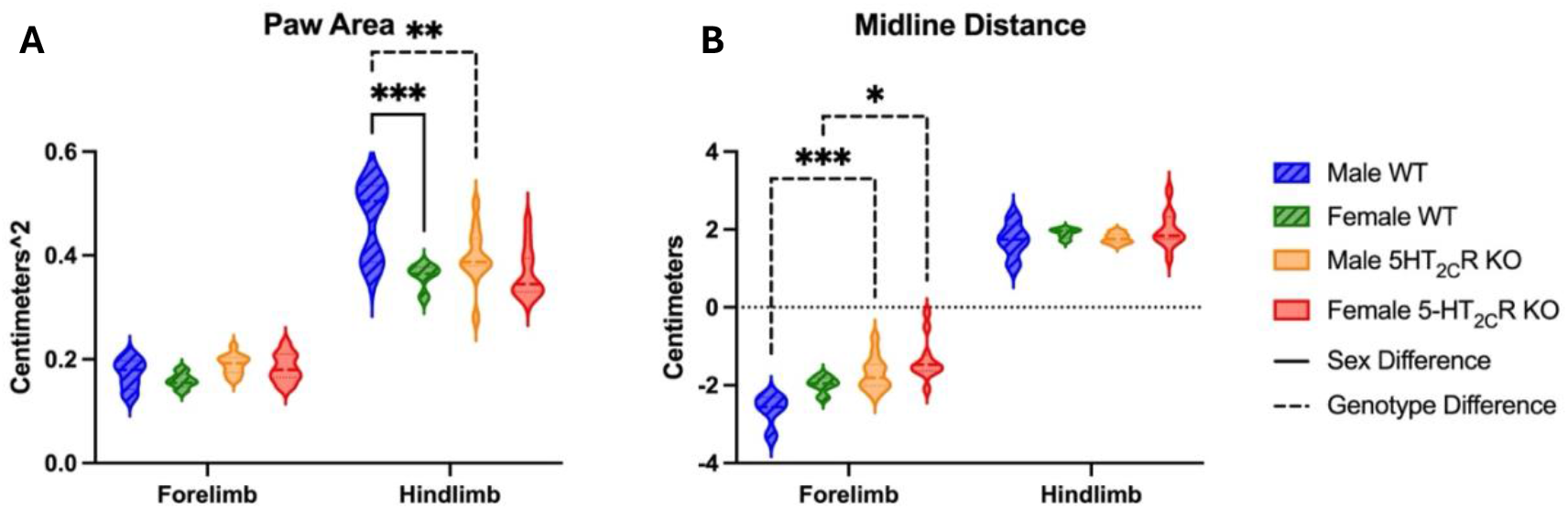
The 5-HT_2C_R KO mice exhibit smaller paw area and shorter midline distance. A two-way ANOVA revealed significant differences in hindlimb paw area and forelimb midline distance. Male WT (n = 5), female WT (n = 5), male 5-HT_2C_R KO (n = 12), and female 5-HT_2C_R KO (n = 11) mice were used in this analysis. A solid significance line indicates a sex-specific difference, and a dashed significance line indicates a genotype-specific difference. **** p < 0.0001, *** p < 0.001, ** p < 0.01, * p < 0.05.

After assessing all gait parameters, we deemed it necessary to investigate whether the size of the mouse had any significant effect on stride length, stance width, and midline distance – all parameters that could be influenced by a longer or wider mouse. Supplementary Table 1 shows the effect of mouse length and width on these specific parameters. Notably, male WT mice were the predominant mouse group affected by mouse length, with forelimb stance width, forelimb midline distance, and hindlimb midline distance all having a significant interaction (*P* = 0.0386, *P* = 0.0297, and *P* = 0.0032, respectively). Additionally, female 5-HT_2C_R KO mice also had a significant interaction between mouse length and forelimb midline distance (*P* = 0.0079). The width of male WT mice, as well as male and female 5-HT_2C_R KO mice had a significant effect on hindlimb stance width (*P* = 0.0004, *P* = 0.0073, and *P* = 0.0498, respectively). Additionally, the width of male WT mice had a significant effect on forelimb midline distance (*P* = 0.0143) and the width of female 5-HT_2C_R KO mice had a significant effect on hindlimb midline distance (*P* = 0.0203). These results suggest that the individual mouse length and width may have a significant effect on specific gait parameters assessed using the DigiGait™ system.

### Female 5-HT_2C_R KO Mice Exhibit Reduced Hindlimb and All-Limb Grip Strength

The differences in gait, paw area, and midline distance reported above suggested changes in motor control, stability, and coordination. Since volitional locomotion is influenced by muscle strength, we investigated the average grip strength in forelimb, hindlimb, and all-limb (Figure 4B). In addition, we evaluated the peak force generated during a single, maximum-effort trial (Figure 4C). In the hindlimb, both average (*P* < 0.001) and maximum (*P* = 0.0093) grip strength was reduced in female 5-HT_2C_R KO mice compared to female WT mice, suggesting that female 5-HT_2C_R KO mice demonstrate overall general muscle weakness compared to their WT counterparts.

**Figure 4.**
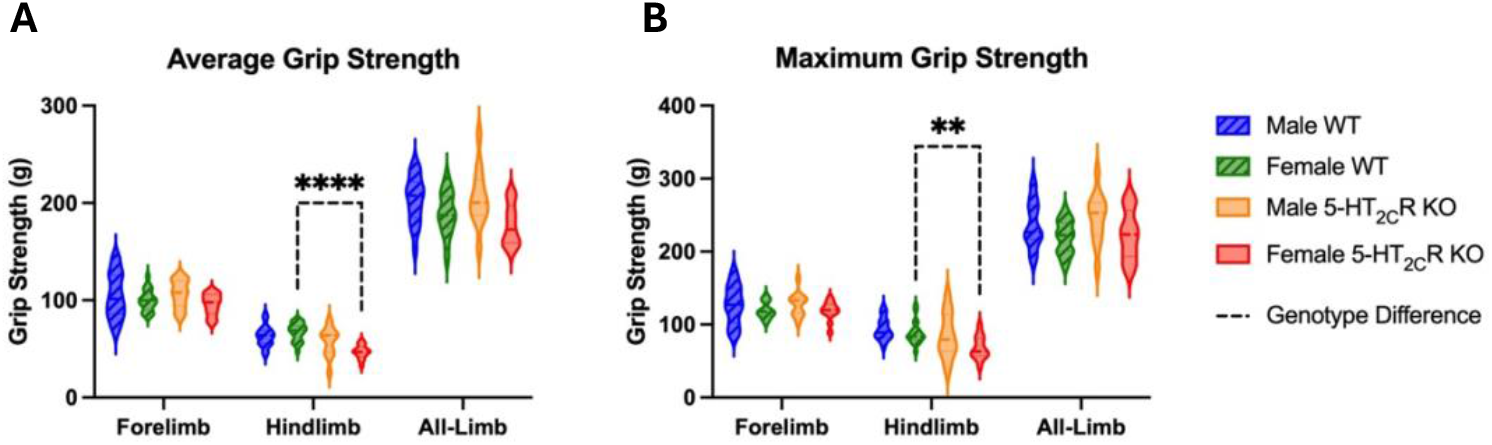
Female 5-HT_2C_R KO mice exhibit decreased muscle weakness compared to female WT mice. A two-way ANOVA revealed significant differences in the average (A) and maximum (B) hindlimb grip strength of female WT and 5-HT_2C_R KO mice. Male WT (n = 18), female WT (n = 18), male 5-HT_2C_R KO (n = 12), and female 5-HT_2C_R KO (n = 12) mice were used in this analysis. A dashed significance line indicates a genotype-specific difference. **** p < 0.0001, *** p < 0.001, ** p < 0.01, * p < 0.05.

### Gross Motor Recovery is Not Affected by 5-HT_2C_ Receptors

To test the effect of the 5-HT_2C_R KO in functional recovery and involuntary motor behaviors, we induced a complete SCI (transection) in both WT and KO mice. An open-field behavioral assessment was performed weekly and scored using the BMS over an 11-week period, starting one day post-SCI (acute) and continuing for 10 weeks (chronic). There were no differences in BMS scores between male and female mice of either genotype, therefore data was combined within each group. By week 10, neither the WT or 5-HT_2C_R KO mouse group scored above a 2 – which is characterized by extensive ankle movement but no coordinated or consistent stepping activity (Figure 5A). Importantly, there was no difference between WT and 5-HT_2C_R KO mice at week 10 (Figure 5B), suggesting that the gross motor recovery of both groups of mice did not differ from one another.

**Figure 5.**
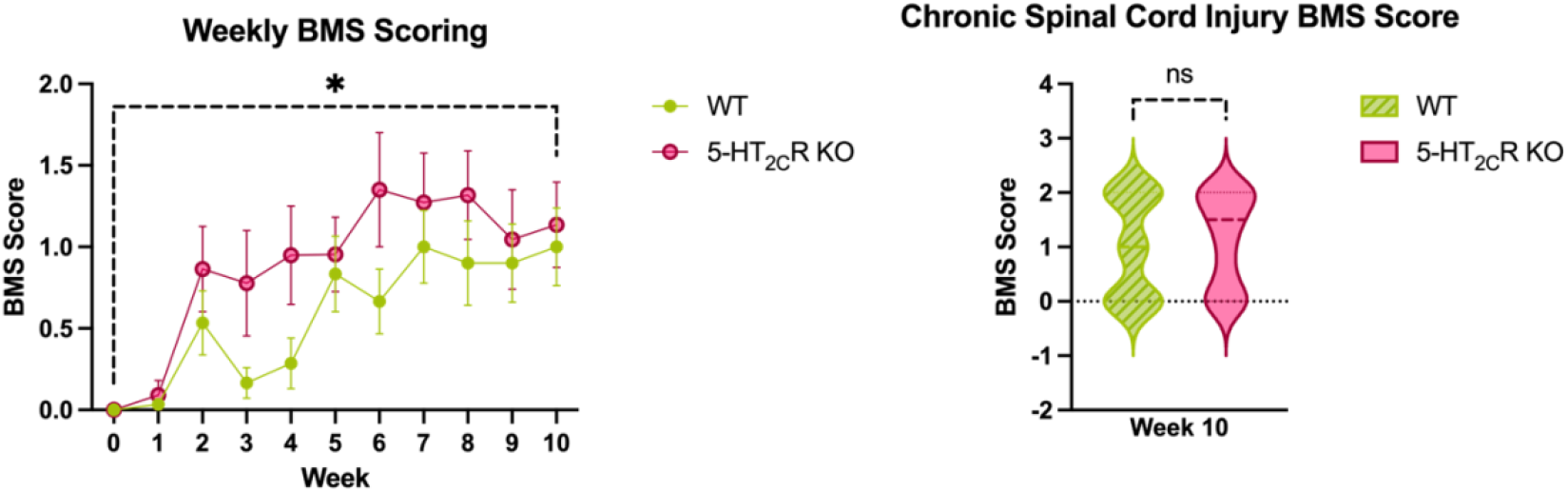
No difference observed in functional motor recovery post-SCI between WT and 5-HT_2C_R KO mice. Scatter plot illustrating (A) the full data set of BMS scoring between WT (n = 15) and 5-HT_2C_R KO mice (n = 11) from week 0 to week 10, and a violin plot illustrating (B) the comparison of BMS scoring between WT (n = 15) and 5-HT_2C_R KO mice (n = 11) on week 10 when the injury had reached chronicity. A two-way ANOVA was used for statistical analysis in (A), and an unpaired t-test was used for statistical comparison in (B). The dashed line on (B) is representative of the median grip strength. **** p < 0.0001, *** p < 0.001, ** p < 0.01, * p < 0.05.

### 5-HT_2C_R KO Mice Exhibited Less Hyperreflexia than WT Mice Post-SCI

To assess whether 5-HT_2C_R KO affects hyperreflexia following SCI, we tested withdrawal reflexes *in vivo*. We electrically stimulated the plantar side of the hind paw while recording EMG activity in the LG and TA antagonistic muscle pair. The evoked reflex was compared as total EMG signal and was broken down into multiple phases (LPR and LLR). There was no difference in LG activity except for female 5-HT_2C_R KO mice having greater total EMG signal than female WT mice (*P* = 0.0311) (Figure 6A). In the TA, however, both male and female 5-HT_2C_R KO mice had lower total EMG signal than their counterpart WT mice (*P* = 0.0098 and 0.0348, respectively; Figure 6A). We then combined male and female data to view non-sex-specific differences in flexor withdrawal caused by the loss of the receptor. The data revealed no significant differences in reflex amplitude in the LG muscle (Figure 6B). However, in the TA, we found lower total EMG signal (*P* = 0.0399) in the KO mice compared to WT (Figure 6B). This indicates that the loss of the functional 5-HT_2C_R resulted in reduced hyperreflexia following SCI.

**Figure 6.**
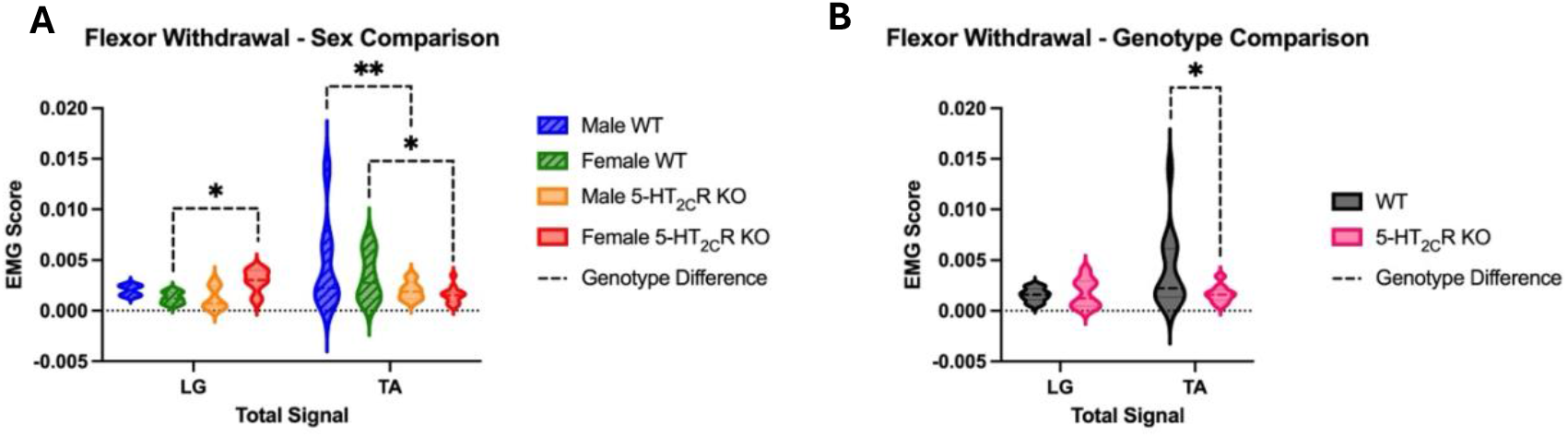
5-HT_2C_R KO mice exhibit decreased hyperreflexia compared to WT mice. Violin-plot illustrating sex-differences between male WT (n = 8), female WT (n = 6), male 5-HT_2C_R KO (n = 5), and female 5-HT_2C_R KO (n = 12) in the LG and the TA (A). Combined male and female data for WT (n = 14) and 5-HT_2C_R KO (n = 17) mice shown in the LG and TA (B). A two-way ANOVA was used for statistical analysis and the dashed significance line is indicative of a genotype-specific difference. **** p < 0.0001, *** p < 0.001, ** p < 0.01, * p < 0.05.

### 5-HT_2C_R KO Mice Exhibited Less Spasm-Like Activity Ex Vivo Post-SCI Compared to WT Mice

The reduction in hyperreflexia observed in 5-HT_2C_R KO mice is likely due to a decrease in MN PICs, which are known to be facilitated by this receptor. However, our recent work has shown that sensory-evoked spastic activity is also influenced by synaptic inhibition in the spinal cord (Mahrous et al., 2024). To test whether changes in inhibition contributed to the observed decrease in hyperreflexia, we measured sensory-evoked spastic activity *ex vivo* in the sacrocaudal spinal cord while blocking synaptic inhibition. We used 2xT and 10xT stimulation of the dorsal roots on either side of the cord to activate low threshold and high threshold afferents, respectively. The evoked ventral root responses were broken down into multiple phases (similar to *in vivo* EMG data described above). In general, male 5-HT_2C_R KO mice showed lower evoked VR activity than male WT mice at different phases of the evoked activity in 2xT contralateral stimulation (LLR *P* < 0.0001, Total Signal *P* < 0.0001; Figure 7A) and 10xT contralateral stimulation (LLR *P* < 0.0001, Total Signal *P* < 0.0001; Figure 7B).

**Figure 7.**
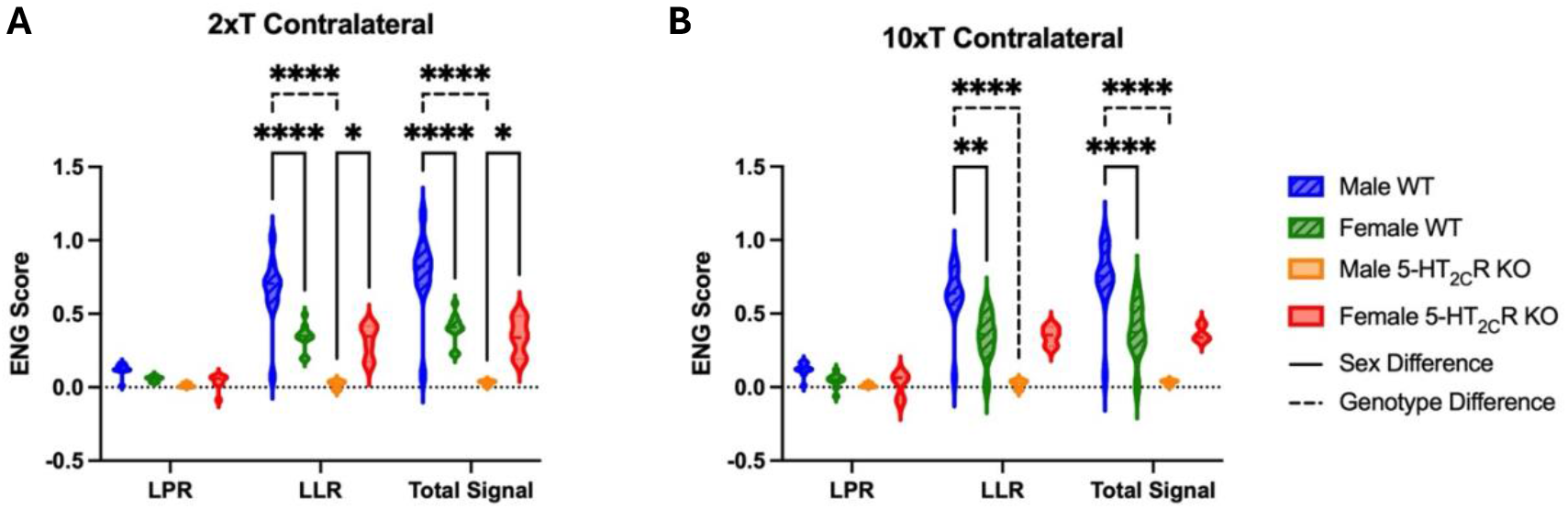
Male 5-HT_2C_R KO mice exhibited less spasm-like activity than male WT mice. Violin plot showing integrated signal results for sacral cord preparation in male WT (n = 6), female WT (n = 9), male 5-HT_2C_R KO (n = 2), and female 5-HT_2C_R KO (n = 5) at 2xT contralateral stimulation (A) and 10xT contralateral stimulation (B). A two-way ANOVA was used for statistical analysis and a solid significance line is indicative of a sex-specific difference and a dashed significance line is indicative of a genotype-specific difference. **** p < 0.0001, *** p < 0.001, ** p < 0.01, * p < 0.05.

Numerous sex-specific differences were revealed in the *in vivo* assessment. Male WT mice had greater ENG signal in LLR and total signal than female WT mice at both 2xT (LLR *P* < 0.0001, total signal *P* < 0.0001; Figure 7A) and 10xT (LLR *P* = 0.0043, total signal *P <* 0.0001; Figure 7B). Conversely, female 5-HT_2C_R KO mice showed greater evoked VR activity than male 5-HT_2C_R KO mice at 2xT in LLR and total signal (*P* = 0.0444 and *P* = 0.0209, respectively; Figure 7A).

These results align with what was observed in the flexor withdrawal reflex assessment, specifically that the 5-HT_2C_R KO mice consistently exhibit lower ENG scores than WT mice, even while synaptic inhibition is blocked. This indicates that the 5-HT_2C_R plays a prominent role in enhancing spasms in chronic SCI, mostly through facilitating MN PICs.

### Both Sex- and Genotype-Specific Differences Revealed in Western Blot Protein Analysis

All three of the 5-HT_2_ receptor subtypes have been shown to possess similar molecular structure, pharmacology, and signal transduction pathways to one another (Parajulee and Kim, 2023). Of the three 5-HT_2_ receptor subtypes, 5-HT_2A_ and 5-HT_2C_ have been the most extensively studied for their role in motor function. It is unclear as to whether the 5-HT_2C_R or 5-HT_2A_R is upregulated post-SCI, but previous research using total protein analysis and mRNA analysis have presented conflicting results. Previous total mRNA studies examining rats that have undergone sacral transection have suggested that the 5-HT_2C_R is not upregulated after sacral transection; however, constitutive activity of the 5-HT_2C_R does become upregulated post-SCI (Murray et al., 2010). Conversely, immunoreactivity studies and total protein analyses of the 5-HT_2C_R have suggested that the 5-HT_2C_R protein is upregulated in the sacral cord 60 days post sacral spinal transection as compared to sham rats (Murray et al., 2010; Ren et al., 2013). In research conducted to investigate the closely related 5-HT_2A_R, it has been suggested that the 5-HT_2A_R is upregulated in the sacral spinal cord post sacral spinal transection (Kong et al., 2010).

To further examine possible mechanistic alterations that occur in WT and 5-HT_2C_R KO mice post-SCI, we used whole spinal cord tissue western blot protein analysis to quantify the relative expression of 5-HT_2C_ and 5-HT_2A_ in both the lumbar and sacral spinal cord of both uninjured and injured WT and 5-HT_2C_R KO mice (Figure 8A). There were no significant differences revealed in the distribution of the 5-HT_2C_ receptor in either the lumbar or sacral spinal cord of male or female WT mice. Conversely, both uninjured female WT mice and uninjured female 5-HT_2C_R KO mice had greater relative expression of the 5-HT_2A_ receptor in the sacral spinal section compared to the lumbar spinal section (data not shown; *P* < 0.0001 and *P* = 0.0004, respectively).

**Figure 8.**
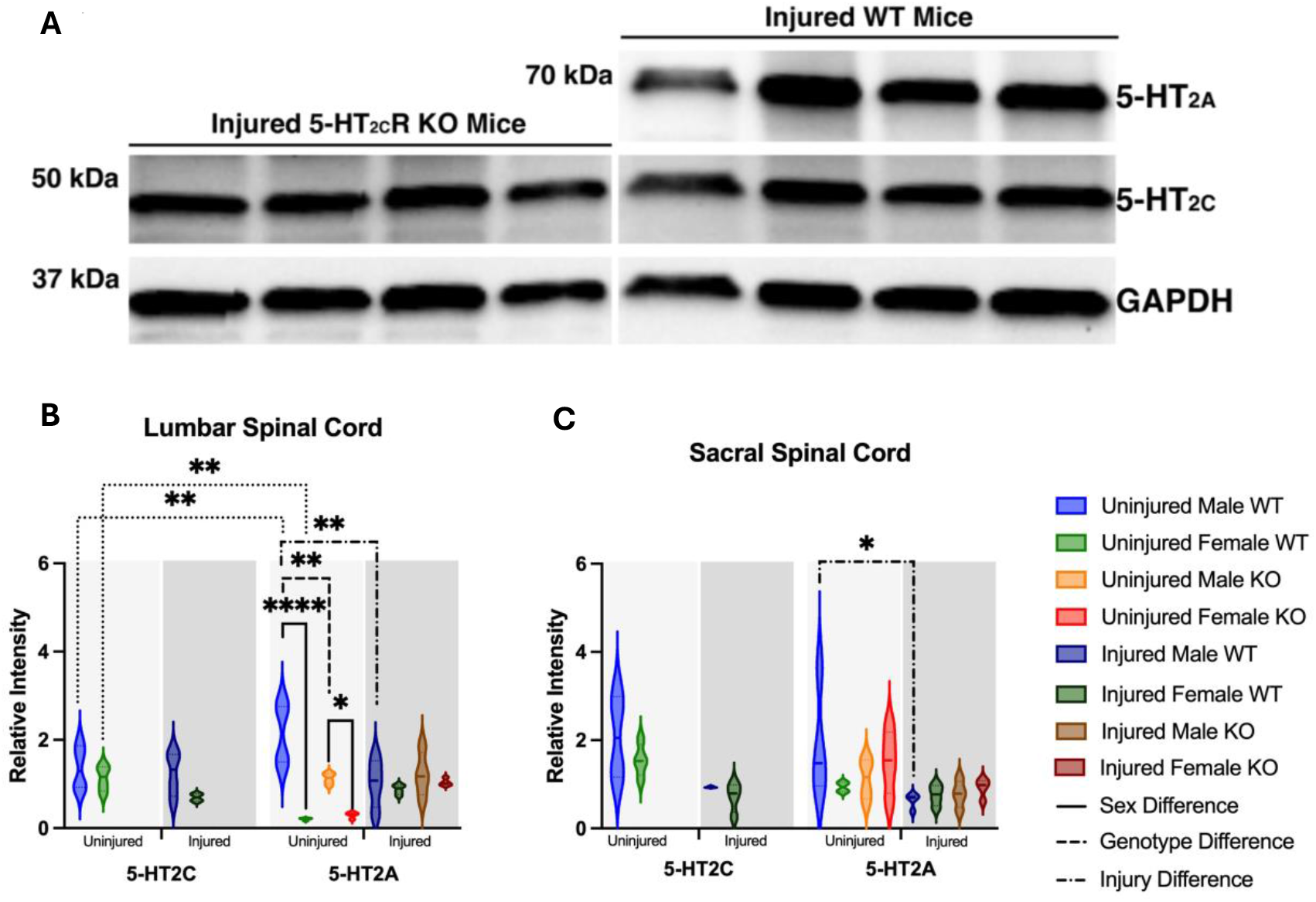
Sex, genotype, and injury status all have an effect on the relative expression and distribution of the 5-HT_**2A**_R. Representative western blot (A) in which the lanes 1 – 4 are the lumbar section of injured, male 5-HT_2C_R KO mice and lanes 5 – 8 are the lumbar section of injured, male WT mice. The relative expression of the 5-HT2CR and 5-HT2AR in the lumbar (B) and sacral (C) spinal cord of uninjured male WT (n = 4), uninjured female WT (n = 4), uninjured male 5-HT_2C_R KO (n = 4), uninjured female 5-HT_2C_R KO (n = 4), injured male WT (n = 4), injured female WT (n = 4), injured male 5-HT_2C_R KO (n = 4), injured female 5-HT_2C_R KO (n = 4) mice. A two-way ANOVA was used for statistical analysis and a solid significance line is indicative of a sex-specific difference, a dashed significance line is indicative of a genotype-specific difference, and a dotted and dashed significance line is indicative of an injury-specific difference. **** p < 0.0001, *** p < 0.001, ** p < 0.01, * p < 0.05.

Comparing the lumbar and sacral spinal sections individually, a multitude of significant differences were identified in regard to 5-HT_2A_ and 5-HT_2C_ receptor expression across sex, genotype, and injury. Figure 8B shows that uninjured male WT mice exhibited significantly greater 5-HT_2A_ receptor relative expression compared to 5-HT_2C_ receptor relative expression in the lumbar spinal cord (*P* = 0.0032), whereas female WT mice displayed the opposite pattern in the same spinal section, with greater 5-HT_2C_ receptor expression relative to 5-HT_2A_ receptor expression (*P* = 0.0005). A similar trend was observed in the lumbar cord of uninjured female 5-HT_2C_R KO mice, where 5-HT_2C_ receptor relative expression exceeded 5-HT_2A_ receptor relative expression (*P* = 0.0384), which may suggest the sex-dependent regulation of serotonergic receptor expression that persists in the absence of the 5-HT_2C_ receptor.

Sex differences were further evident in 5-HT_2A_ receptor relative expression, as uninjured male WT mice had significantly greater 5-HT_2A_ receptor relative expression compared to uninjured female WT mice in the lumbar spinal cord (*P* < 0.0001; Figure 8B). This same trend was also observed in the 5-HT_2C_R KO mice, with uninjured male 5-HT_2C_R KO mice having greater 5-HT_2A_ receptor relative expression than uninjured female 5-HT_2C_R KO mice in the lumbar spinal cord (*P* = 0.0345; Figure 8B), further suggesting that the sex-specific regulation of 5-HT_2A_ receptor is independent of 5-HT_2C_ receptor presence.

A significant genotype-specific effect was also observed, as uninjured male WT mice exhibited greater 5-HT_2A_ receptor relative expression compared to uninjured male 5-HT_2C_R KO mice in the lumbar spinal cord (*P* = 0.0043; Figure 8B), indicating that 5-HT_2C_ receptor absence may influence 5-HT_2A_ receptor regulation in uninjured mice.

Injury-induced changes were also evident in male WT mice, as 5-HT_2A_ receptor relative expression was significantly lower in injured compared to uninjured male WT mice in both the lumbar and sacral spinal cord (*P* = 0.0012 and *P* = 0.0311, respectively; Figure 8B – C), possibly suggesting that spinal cord injury may lead to a downregulation of 5-HT_2A_ receptor expression across multiple spinal regions.

## Discussion

This study was conducted to gain a more comprehensive understanding of how the 5-HT_2C_R is implicated in volitional and involuntary motor behavior in uninjured and SCI adult mice. The main findings of this study are as follows: 1) the absence of the 5-HT_2C_R is associated with alterations in volitional locomotion and muscular strength; 2) the 5-HT_2C_R has an effect on the severity and prevalence of involuntary motor behavior post-SCI; and 3) the relative expression of the 5-HT_2C_R and 5-HT_2A_R is influenced by sex, genotype, and injury-status of the mouse.

### 5-HT_2C_R KO mice lack the strength and stability required for normal locomotion

Examining normal motor control via highly-specified gait analysis and grip strength assessment using a receptor knock-out model can provide valuable insight into receptor function. Previous research has shown varying involvement and implication of the 5-HT_2C_R in motor functioning in healthy mice, yet to our best knowledge, there has been no study that has examined potential sex differences. In this study, we used the DigiGait™ analysis alongside a comprehensive grip strength assessment which provided information concerning locomotion and muscular strength, respectively, of both WT and 5-HT_2C_R KO mice. It was hypothesized that the 5-HT_2C_R KO mice would perform worse on the volitional motor control assessments – as the link between the 5-HT_2C_R and spinal locomotion has been previously established (Majczyński et al., 2020).

There were numerous differences observed in the DigiGait™ digital treadmill gait analysis that supported our hypothesis. Notably, the male 5-HT_2C_R KO mice spent an increased amount of time in stance and a decreased amount of time in swing compared to the WT mice in the forelimb analysis, and the female 5-HT_2C_R KO and WT mice exhibited the same trend in the hindlimb. It is important to note that stance is a double legged position, whereas swing is a single legged position. The observation that 5-HT_2C_R KO mice spent a greater amount of time in stance but not in swing suggests that these mice lack the stability and power found in their WT counterparts. Furthermore, when the stance phase of gait was divided into brake and propel, male 5-HT_2C_R KO mice spent an increased amount of the stance phase in brake and a decreased amount in propel compared to male WT mice. This suggests that the male 5-HT_2C_R KO mice are unable to efficiently balance on the toe of the paw and instead tend to spend a greater percentage of the stance phase with the heel of the paw on the treadmill belt – a more stable position that enables the mouse to remain balanced while walking.

In addition to having significant differences in gait phase composition, we also hypothesized that the 5-HT_2C_R KO mice would exhibit overall poorer strength, possibly resulting in deficits such as decreased posturing ability, compared to the WT mice. This hypothesis was supported by the results obtained from the paw area and midline distance analysis. The male 5-HT_2C_R KO mice had a smaller hindlimb paw area than male WT mice, suggesting that the strength required to splay the hindlimb paw was decreased. This may contribute to the differences observed in the breakdown of the stance phase of gait in which the male 5-HT_2C_R KO mice had a tendency to remain on the heel of the paw instead of the toes of the paw. As for the midline distance analysis, both male and female 5-HT_2C_R KO mice exhibited a narrower forelimb midline distance than male and female WT mice, respectively. This suggests that the 5-HT_2C_R KO mice possess an inability to fully extend the forelimb outward from the midline. A narrower stance is inherently more unstable than a wider stance, supporting the hypothesis that the 5-HT_2C_R KO mice lack adequate stability and gait posturing that WT mice naturally exhibit.

After determining that there were numerous significant differences in gait and locomotion between the 5-HT_2C_R KO mice and WT mice, we furthered our examination into volitional behavior of the two mouse groups by assessing the average and maximum grip strength of the mice. We hypothesized that the 5-HT_2C_R KO mice would have decreased grip strength based on what had been observed in the DigiGait™ analysis, and this hypothesis was supported in the female but not male mice. Average and maximum hindlimb grip strength was significantly lower in female 5-HT_2C_R KO mice compared to female WT mice. In addition to this genotype-specific result, female WT mice had lower average and maximum hindlimb grip strength than male WT mice, and a similar trend was seen in the all-limb average grip strength result as well. Previous research has suggested that female mice have larger PICs in lower limb MNs than male mice (Jenz et al., 2023), which may contribute to the difference in grip strength being observed in only the female 5-HT_2C_R KO and WT mice.

### 5-HT_2C_R KO mice exhibit decreased involuntary motor behavior

Following the examination of volitional motor behavior in uninjured 5-HT_2C_R KO and WT mice, an SCI model was implemented to evaluate possible differences in involuntary motor behavior between the two groups of mice. We hypothesized that the injured 5-HT_2C_R KO mice would exhibit decreased involuntary motor behaviors compared to the WT mice because there would be no constitutive activity of the 5-HT_2C_R below the level of injury, which has been previously suggested to contribute to MN PIC activity and spasms in SCI mice (Murray et al., 2011a). Our findings show that the 5-HT_2C_R KO mice had consistently lower EMG scores in each LLR phase than WT mice, suggesting that the 5-HT_2C_R KO mice exhibited overall lower reflex hyperexcitability post-SCI compared to WT mice. The discrepancy in the presentation of involuntary motor behavior is likely related to the absence of constitutive activity – which requires the functional 5-HT_2C_R to contribute to the generation of PIC’s, thereby eliciting spasm-like activity.

Neural activity associated with spasm-like activity in SCI mice was further evaluated by conducting *ex vivo* whole-tissue sacral cord preparation. By implementing this *ex vivo* experimental technique, confounding variables such as mouse movement and environmental stimuli – both of which are possible limitations of flexor withdrawal response testing – were eliminated. Synaptic inhibition was blocked using STR/PTX to inhibit interneuron activity, thereby increasing the length of the monosynaptic response to measure the LLR response.

Previous research has shown that consecutive dorsal root stimulation at 2 times threshold (see methods) every 15 s will cause evoked spasm-like activity to adapt, decreasing in duration until reaching a plateau after 8 to 10 spasms (Mahrous et al., 2024). Findings reveal that 5-HT_2C_R KO mice had decreased ENG scores for both the ipsilateral and contralateral ventral root monosynaptic response compared to WT mice – a result that aligns with the results of the flexor withdrawal analysis. Importantly, the finding that 5-HT_2C_R KO mice having decreased ENG score compared to WT mice was predominately shown in male mice. However, female 5-HT_2C_R KO mice also had lower ENG score compared to female WT mice at 10xT on the ipsilateral side in LLR5 and total signal. This finding suggests that the 5-HT_2C_R does have a critical role in the contribution to spasm-like activity following SCI.

### Sex, Genotype, and Injury-Dependent Modulation of 5-HT_2C_R and 5-HT_2A_R Expression in the Lumbar and Sacral Spinal Cord

A western blot protein analysis using whole-tissue spinal cord was conducted following behavioral and *ex vivo* experiments. This approach aimed to investigate the underlying molecular mechanisms that may explain the differences observed in previous behavioral experiments. By quantifying the relative expression of the 5-HT_2C_R and 5-HT_2A_R in the lumbar and sacral spinal cord, we sought to determine whether alterations in serotonergic signaling contribute to the sex, genotype, and injury-dependent differences in motor behavior.

We initially hypothesized that the absence of the 5-HT_2C_R would lead to an up-regulation of the 5-HT_2A_R through a compensatory molecular mechanism, but this was not supported. Instead, uninjured male 5-HT_2C_R KO mice exhibited lower 5-HT_2A_R relative expression in the lumbar spinal cord compared to uninjured male WT mice, suggesting that the absence of the 5-HT_2C_R does not drive increased 5-HT_2A_R expression. Previous research has suggested that a chronic blockade of the 5-HT_2C_R and 5-HT_2A_R results in down-regulation rather than the expected up-regulation (Van Oekelen et al., 2003), which corroborates our findings here.

Previous research has suggested that the 5-HT_2A_R becomes upregulated in male Wistar rats after sacral transection (Kong et al., 2011, p. 22, 2010), so we initially hypothesized that injured mice would exhibit increased relative expression of the 5-HT_2A_R compared to uninjured mice – regardless of sex or genotype. Our findings revealed the opposite; injured male WT mice had *lower* relative expression of the 5-HT_2A_R in both the lumbar and sacral spinal cord compared to uninjured male WT mice. It is important to note that the studies cited above used immunofluorescence, whereas our study used whole tissue western blot protein analysis.

Additionally, the studies above employed a sacral transection on male rats, whereas our study used the more clinically relevant thoracic transection on mice of both sexes.

There was one significant regional difference revealed in 5-HT_2A_R relative expression between the lumbar and sacral spinal cord. Both uninjured female WT and uninjured female 5-HT_2C_R KO mice had greater 5-HT_2A_R relative expression in the sacral spinal cord compared to the lumbar spinal cord. This finding suggests that there is a notable difference in 5-HT_2A_R distribution in uninjured mice, regardless of the presence of the 5-HT_2C_R, but only in female mice. Additionally, both uninjured female WT and uninjured female 5-HT_2C_R KO mice exhibited lower 5-HT_2A_R relative expression than uninjured male WT and uninjured male 5-HT_2C_R KO mice in the lumbar spinal cord. So not only did female mice (both WT and 5-HT_2C_R KO) have a greater distribution of relative expression of the 5-HT_2A_R in the sacral spinal cord, but they also exhibited decreased 5-HT_2A_R relative expression in the lumbar spinal cord compared to male WT and 5-HT_2C_R KO mice. These results suggest there may be a sex hormone-driven regulation of 5-HT_2A_R expression. Little to no research has been conducted investigating how sex hormones affect the serotonergic system in the spinal cord, but there have been many studies that have examined the role of estradiol and testosterone and their respective effects on specific serotonergic receptors within the brain (Kugaya et al., 2003; Sumner and Fink, 1998). Our results suggest that there may be a connection between sex-specific and hormonal differences in 5-HT_2A_R and 5-HT_2C_R relative expression in the spinal cord of uninjured and SCI mice, regardless of whether the 5-HT_2C_R is functional or not.

## Limitations

A limitation of this study is the lack of EMG normalization in signal processing for the flexor withdrawal analysis. In human studies, EMG signal analysis is typically normalized to the maximum volitional muscle contraction of the subject. In this study, as well as in other animal studies, initiating a voluntary maximum muscle contraction within each subject is difficult and may impact the signal-to-background noise and subsequent calculations.

Additionally, the menstrual cycle of female mice was not tracked. This is a limitation of the study because potential fluctuations in hormones that have been previously shown to impact the serotonergic system were not accounted for in our analyses. The menstrual cycle of female mice may have a significant impact on 5-HT_2A_R and 5-HT_2C_R expression, directly impacting the results of our western blot analyses and possibly extending outward to affect our volitional behavioral testing as well. Future studies investigating the effect of serotonergic receptors on male and female mouse behavior and tissue must consider tracking the menstrual cycle of female mice to provide further clarity on hormonal alterations that may directly impact their results.

### Future Directions

This study provides comprehensive behavioral insight and immunohistochemical analysis on how the absence of the functional 5-HT_2C_R can affect volitional and involuntary motor behavior, and how the expression of the 5-HT_2A_R is altered in the sacral and lumbar spinal cord post-SCI. This study only examined two specific serotonergic receptors (5-HT_2A_R and 5-HT_2C_R) and their possible effect on MN excitability post-SCI. However, serotonergic input is not the only input that is capable of mediating MN excitability post-SCI. The noradrenaline receptor, for example, has been shown to facilitate the recovery of PICs following a SCI (Heckman et al., 2009). Future research might consider additional analysis tailed to compare serotonergic receptors and norepinephrine receptors in order to determine whether one source of input has a larger effect on involuntary motor behavior post-SCI.

Previous research has suggested that spinalized rats develop increased immunoreactivity of the 5-HT_2A_R below the level of lesion in the sacrocaudal spinal cord. The data from our study does not support this, when comparing the expression of the 5-HT_2A_R in the sacral cord of injured to uninjured WT mice. However, it is not fully understood whether the supersensitivity of the 5-HT_2A_R below the level of lesion is a result of conformational changes in the receptor itself, or by changes in cellular signaling. Understanding that PICs are facilitated by the presence of monoamines such as 5-HT, it could be plausible that the 5-HT_2A_R undergoes changes in constitutive activity similar to those identified in the 5-HT_2C_R. Previous electrophysiology studies have suggested that the 5-HT_2A_R has a significant effect on MN excitability in the hindlimb, but it is widely understood that these changes are not solely due to the 5-HT_2A_R itself (Majczyński et al., 2020). Future studies might consider incorporating additional immunohistochemical analyses, such as immunofluorescence staining, to better understand changes in 5-HT_2A_R and 5-HT_2C_R localization post-SCI, specifically in the plasma membrane and cytoplasm of the MN.

Gaining a comprehensive understanding as to how the 5-HT_2C_R and 5-HT_2A_R are involved in both volitional and involuntary movement post-SCI is necessary in order to determine how these receptors support either volitional recovery and/or severity of hyperreflexia/spasms. Examining how the expression and quantity of the 5-HT_2C_R and 5-HT_2A_R is altered in the lumbar and sacral spinal cord post-SCI may provide further insight that would allow for the development of novel and precise therapeutic strategies that target these specific to assist in reducing the frequency and severity of muscle spasms post-SCI.

