## Supplemental Table 1 for "Loss of 5-HT_2C_ Receptor Function Alters Motor Behavior in Male and Female Mice With and Without Spinal Cord Injury"

### Supplementary Material

**Table 1.** Statistical correlation between mouse length/width and DigiGait™ parameters.

| | Comparison | Correlation (R) | P-Value | n | 95% CI | Significance |
|---|---|---|---|---|---|---|
| Forelimb | Male WT Mouse Length vs. Stride Length | -0.4272 | 0.4731 | 5 | -0.9510 to 0.7303 | ns |
| | Female WT Mouse Length vs. Stride Length | -0.6374 | 0.2474 | 5 | -0.9727 to 0.5595 | ns |
| | Male 5-HT2CR KO Mouse Length vs. Stride Length | 0.4193 | 0.1748 | 12 | -0.2036 to 0.8006 | ns |
| | Female 5-HT2CR KO Mouse Length vs. Stride Length | 0.2881 | 0.3903 | 11 | -0.3769 to 0.7571 | ns |
| Hindlimb | Male WT Mouse Length vs. Stride Length | 0.1593 | 0.798 | 5 | -0.8412 to 0.9132 | ns |
| | Female WT Mouse Length vs. Stride Length | -0.6159 | 0.2687 | 5 | -0.9707 to 0.5834 | ns |
| | Male 5-HT2CR KO Mouse Length vs. Stride Length | 0.4742 | 0.1194 | 12 | -0.1370 to 0.8239 | ns |
| | Female 5-HT2CR KO Mouse Length vs. Stride Length | 0.01227 | 0.9714 | 11 | -0.5920 to 0.6077 | ns |
| Forelimb | Male WT Mouse Length vs. Stance Width | 0.8977 | 0.0386 | 5 | 0.07442 to 0.9933 | * |
| | Female WT Mouse Length vs. Stance Width | -0.006013 | 0.9923 | 5 | -0.8836 to 0.8809 | ns |
| | Male 5-HT2CR KO Mouse Length vs. Stance Width | -0.01902 | 0.9532 | 12 | -0.5865 to 0.5610 | ns |
| | Female 5-HT2CR KO Mouse Length vs. Stance Width | -0.1365 | 0.6889 | 11 | -0.6807 to 0.5047 | ns |
| Hindlimb | Male WT Mouse Length vs. Stance Width | 0.8247 | 0.0857 | 5 | -0.2112 to 0.9881 | ns |
| | Female WT Mouse Length vs. Stance Width | 0.3263 | 0.5921 | 5 | -0.7807 to 0.9384 | ns |
| | Male 5-HT2CR KO Mouse Length vs. Stance Width | -0.06415 | 0.843 | 12 | -0.6154 to 0.5292 | ns |
| | Female 5-HT2CR KO Mouse Length vs. Stance Width | -0.004231 | 0.9901 | 11 | -0.6026 to 0.5972 | ns |
| Forelimb | Male WT Mouse Length vs. Midline Distance | -0.9143 | 0.0297 | 5 | -0.9944 to -0.1656 | * |
| | Female WT Mouse Length vs. Midline Distance | -0.624 | 0.2606 | 5 | -0.9714 to 0.5747 | ns |
| | Male 5-HT2CR KO Mouse Length vs. Midline Distance | -0.5287 | 0.0772 | 12 | -0.8459 to 0.06484 | ns |
| | Female 5-HT2CR KO Mouse Length vs. Midline Distance | -0.7497 | 0.0079 | 11 | -0.9309 to -0.2722 | ** |
| Hindlimb | Male WT Mouse Length vs. Midline Distance | 0.9807 | 0.0032 | 5 | 0.7307 to 0.9988 | ** |
| | Female WT Mouse Length vs. Midline Distance | 0.5081 | 0.3821 | 5 | -0.6782 to 0.9600 | ns |
| | Male 5-HT2CR KO Mouse Length vs. Midline Distance | 0.4569 | 0.1354 | 12 | -0.1586 to 0.8166 | ns |
| | Female 5-HT2CR KO Mouse Length vs. Midline Distance | -0.4608 | 0.1538 | 11 | -0.8310 to 0.1922 | ns |
| Forelimb | Male WT Mouse Width vs. Stride Length | -0.4173 | 0.4845 | 5 | -0.9499 to 0.7359 | ns |
| | Female WT Mouse Width vs. Stride Length | -0.4042 | 0.4998 | 5 | -0.9483 to 0.7431 | ns |
| | Male 5-HT2CR KO Mouse Width vs. Stride Length | -0.3042 | 0.3363 | 12 | -0.7476 to 0.3267 | ns |
| | Female 5-HT2CR KO Mouse Width vs. Stride Length | 0.3657 | 0.2687 | 11 | -0.3000 to 0.7919 | ns |
| Hindlimb | Male WT Mouse Width vs. Stride Length | -0.3189 | 0.6009 | 5 | -0.9374 to 0.7839 | ns |
| | Female WT Mouse Width vs. Stride Length | 1.205E-15 | >0.9999 | 5 | -0.8823 to 0.8823 | ns |
| | Male 5-HT2CR KO Mouse Width vs. Stride Length | -0.545 | 0.0669 | 12 | -0.8523 to 0.04201 | ns |
| | Female 5-HT2CR KO Mouse Width vs. Stride Length | 0.2832 | 0.3987 | 11 | -0.3814 to 0.7549 | ns |
| Forelimb | Male WT Mouse Width vs. Stance Width | 0.8237 | 0.0865 | 5 | -0.2142 to 0.9880 | ns |
| | Female WT Mouse Width vs. Stance Width | 0.03232 | 0.9589 | 5 | -0.8749 to 0.8892 | ns |

| | | | | | | |
|---|---|---|---|---|---|---|
| | Male 5-HT2CR KO Mouse Width vs. Stance Width | 0.05785 | 0.8583 | 12 | -0.5338 to 0.6115 | ns |
| | Female 5-HT2CR KO Mouse Width vs. Stance Width | 0.1317 | 0.6996 | 11 | -0.5084 to 0.6780 | ns |
| Hindlimb | Male WT Mouse Width vs. Stance Width | 0.9949 | 0.0004 | 5 | 0.9217 to 0.9997 | *** |
| | Female WT Mouse Width vs. Stance Width | -0.1461 | 0.8146 | 5 | -0.9110 to 0.8451 | ns |
| | Male 5-HT2CR KO Mouse Width vs. Stance Width | 0.7277 | 0.0073 | 12 | 0.2640 to 0.9181 | ** |
| | Female 5-HT2CR KO Mouse Width vs. Stance Width | 0.6025 | 0.0498 | 11 | 0.004189 to 0.8832 | * |
| Forelimb | Male WT Mouse Width vs. Midline Distance | -0.9475 | 0.0143 | 5 | -0.9966 to -0.3980 | * |
| | Female WT Mouse Width vs. Midline Distance | 0.2666 | 0.6646 | 5 | -0.8050 to 0.9301 | ns |
| | Male 5-HT2CR KO Mouse Width vs. Midline Distance | 0.03082 | 0.9243 | 12 | -0.5529 to 0.5942 | ns |
| | Female 5-HT2CR KO Mouse Width vs. Midline Distance | 0.1187 | 0.7281 | 11 | -0.5180 to 0.6708 | ns |
| Hindlimb | Male WT Mouse Width vs. Midline Distance | 0.6401 | 0.2448 | 5 | -0.5564 to 0.9729 | ns |
| | Female WT Mouse Width vs. Midline Distance | -0.8064 | 0.0993 | 5 | -0.9867 to 0.2630 | ns |
| | Male 5-HT2CR KO Mouse Width vs. Midline Distance | -0.4556 | 0.1367 | 12 | -0.8161 to 0.1602 | ns |
| | Female 5-HT2CR KO Mouse Width vs. Midline Distance | 0.6838 | 0.0203 | 11 | 0.1424 to 0.9103 | * |
